# Compressive sensing of neuronal connectivity maps from subsampled, cell-targeted optogenetic stimulation of a network model

**DOI:** 10.1101/2022.11.08.515672

**Authors:** Phillip Navarro, Karim Oweiss

## Abstract

Mapping functional connectivity between neurons is an essential step towards probing the neural computations mediating behavior. The ability to consistently and robustly determine synaptic connectivity maps in large populations of interconnected neurons is a significant challenge in terms of yield, accuracy and experimental time. Here we developed a compressive sensing approach to reconstruct synaptic connectivity maps based on random two photon (2p) cell-targeted optogenetic stimulation and membrane voltage readout of many putative postsynaptic neurons. Using a biophysical network model of interconnected populations of excitatory and inhibitory neurons, we found that the mapping can be achieved with far fewer measurements than the standard pairwise sequential approach. We characterized the recall and precision probabilities as a function of network observability, sparsity, number of neurons stimulated per trial, off-target stimulation, synaptic reliability, propagation latency and network topology. We found that that network sparsity and synaptic reliability were primary determinants of the performance. In particular, in a network with 10% probability of neuronal connectivity, functional connections were recovered with >85% recall and >80% precision in half the trials that would be required for single cell stimulation. Our results suggest a rapid and efficient method to reconstruct functional connectivity of brain networks where sparsity is predominantly present.

## Introduction

There is no doubt that single neurons compute, for example, via specific spatiotemporal axosomatic and axodendritic integration^1–10^. It is widely believed, however, that for the brain to support reliable perception and effective action, rapid coordination among ensembles of neurons with heterogenous cell types, morphologies and tuning properties is needed^3,11^. This coordination is enabled by highly precise and dynamic synaptic connectivity maps that vary in size, location and architecture across different brain regions. To map this connectivity, postsynaptic current responses to depolarizing current pulses delivered to presynaptic terminals must be measured using whole cell recordings–a technically challenging technique with very low yield and recording instability, particularly *in vivo*. While two-photon (2p) or robotic-guided whole cell patching^12^ may facilitate this approach, the technique remains extremely slow and cannot be performed repeatedly within a session or across multiple sessions^13^. This precludes the ability to infer large scale synaptic connectivity maps, for example, to characterize neural circuit function subserving a specific behavior^14^ or to track synaptic plasticity associated with learning new behavior^15^.

Recent advances in 2p optogenetics have enabled precise spatial and temporal control of neural activity with single cell resolution in awake behaving animals^16,17^. New high-fidelity opsins with improved kinetic properties coupled with advances in spatial light modulation have made it possible to perform multi-cell stimulation with sub-millisecond precision^18^. In parallel, advances in genetically encoded voltage indicators (GEVIs) have enabled imaging subthreshold membrane potentials in multiple neurons with high enough Signal to Noise Ratios (SNRs)^19–22^. Together, this all-optical toolkit has paved the way for building high resolution synaptic connectivity maps of neuronal circuits in awake behaving animals. Despite these striking advances, the ability to map synaptic connectivity in a large population in a reasonable experimental time scale is still out of reach as neurons have to be stimulated sequentially and independently^23^, rendering most trials with no recorded membrane response ^24–26^ in brain areas known to be sparsely connected such as the neocortex ^27–32^, or in areas where synapses are weak or unreliable ^33^.

In this paper, we demonstrate a rapid compressed connectivity mapping (CoCoMap) approach that uses parallel random stimulation while measuring postsynaptic membrane potential (PSP) responses from multiple cells simultaneously to reconstruct synaptic connectivity parameters (strength and direction). In contrast to the sequential approach, this parallel stimulation approach leverages inferred cell-type information, network sparsity and the theory of compressive sensing (CS)^34,35^ to discover synaptic connectivity with far fewer measurements. Compared to other related work^36–38^, our approach is particularly novel in recapitulating much more biologically plausible network topologies and dynamics such as recurrent connectivity, varying levels of background noise and memory retention of past inputs. We demonstrate that at 10% sparsity typically found in cortical networks^27–32^, CoCoMap achieved >90% performance with only half the number of measurements that would be needed using the sequential approach. Systematically varying model parameters revealed that CoCoMap outperformed the sequential approach over nearly all examined input and network parameter ranges. CoCoMap performance remained particularly robust in the presence of off-target photostimulation effects and highly clustered small-world network architectures. Observing smaller proportions of the underlying network moderately lowered performance, still resulting in >80% performance with half the measurements needed when only 10% of the population in the network could be observed. Performance plateaued as the fraction of cells in a stimulated pattern increased beyond roughly 20% of the observed population, highlighting trade-offs between stimulating a cell and being able to measure its response in the same trial. High synaptic failure probability dramatically lowered performance for a fixed number of measurements, suggesting limitations on the approach’s speed in mapping networks in brain areas known to have unreliable synapses.

## Methods

### Compressive Sensing Background

CS theory is based on the principle that input source signals are sparse in some domain (time, frequency, wavelet, etc.) and that the measurements collected to estimate these source signals are incoherent. Such incoherence can be achieved by parallel stimulation of a random subset of neurons in an imageable population of *N* neurons to generate T measurements of membrane responses (Figure 1A). We define the T x N stimulation matrix M as binary and random where in each of the T rows (T < N) a neuron is stimulated if its respective index equals ‘1’ and not stimulated otherwise. Running this experiment yields a set of 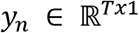 response measurements for each putative postsynaptic neuron *n*, *n=1:N*. We let 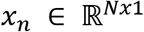 represent the unknown weight vector connecting the stimulated neurons to postsynaptic neuron *n*. Let *e_n_* be the sum of the spontaneous firing *e_n_sf__* and the membrane noise *e_n_ν__* of the postsynaptic neuron. Assume that synaptic currents from stimulated presynaptic neurons are linearly summed, our measured response can be expressed as:

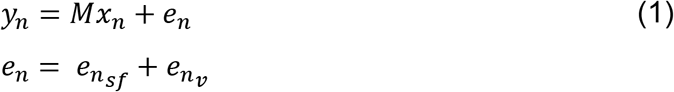

*x* can be estimated from this model by solving

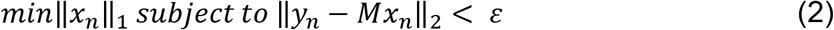

where ||*e_n_*||_2_ ≤ *ε* is an upper bound on the noise. For a sparse signal and incoherent measurements, which the binary random structure of M satisfies^34^, the L1 norm minimization of x subject to the reconstruction error constraint above yields the fewest number of nonzero pre-synaptic weights^39^. This approach, known as basis pursuit^40^, scales relative to the number of connections per neuron as opposed to single input testing which scales relative to the total number of neurons in the network (Figure 1A).

**Figure 1.**
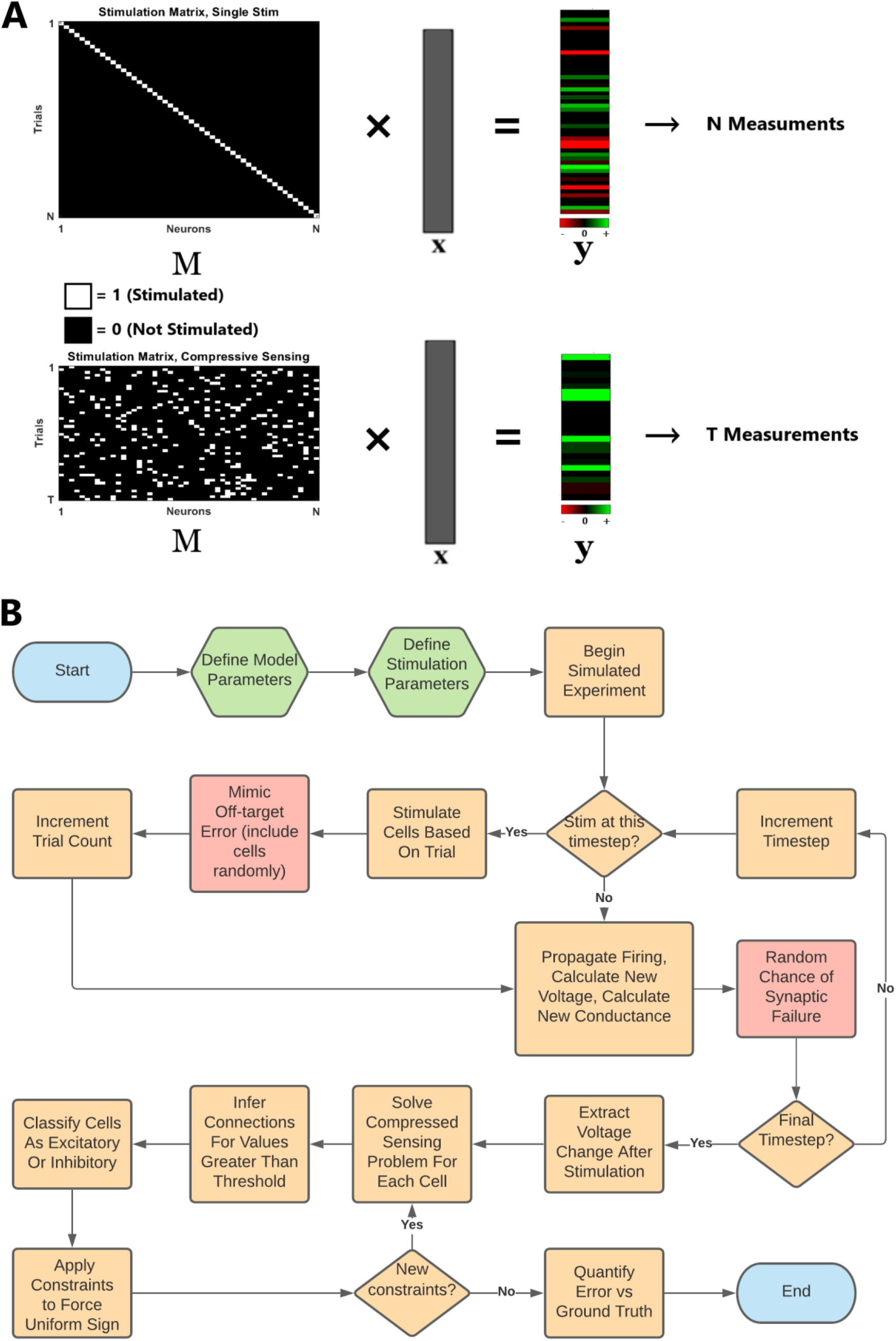
Compressive Connectivity Mapping. A. Measurement strategies for a system of N interconnected sources. M represents a stimulation matrix delivered to N neurons in the network on each of T trials where a measurement y is made of the recorded neuron in each trial x is a vector of the pre-synaptic connection weights to the recorded neuron that we wish to estimate. Top: SISO approach based on perturbing one cell in each trial. This takes a number of measurements equal to the number of neurons with the estimated weights being directly proportional to the measured responses. Bottom: Compressive Connectivity Mapping () applied to a MIMO measurement model. Perturbing multiple neurons per trial then decoding the mixed responses should enable mapping functional connectivity in fewer measurements. B. Simulated Experiment Flow Chart. Orange boxes indicates elements that are part of the base experiment. Red boxes indicate steps that only execute if the respective parameter is non-zero. Green boxes indicate parameter selection and Blue boxes denote the beginning and end of an experiment run. After the model and stimulation experiment parameters are defined, the experiment proceeds by evaluating the differential equations governing the voltage and membrane recovery for each neuron at every timestep. At preset intervals, a stimulation trial is carried out which consists of stimulating neurons corresponding to the indices of the trial row in the binary random matrix M. After the final timestep, changes in voltage corresponding to the timestep after a subset of neurons was stimulated are extracted. These are used to solve the basis pursuit compressive sensing problem (2). Values of x greater than some minimum threshold are said to be connections and compared to the ground truth. Cells are classified into inhibitory and excitatory based on the sign and magnitude of the inferred connections. Projections from these cells are constrained to all be of uniform sign. Error is quantified through metrics of recall and precision.

### Computational Model

In order to test this framework, we developed an in-silico network model composed of 1000 Izhikevich model neurons^41^. The Izhikevich model was chosen for its good balance between biological plausibility, diversity of firing characteristics and computational simplicity. A detailed description of the neuron model is provided in the supplemental materials. For this model, responses are measured as changes in postsynaptic membrane voltages (PSPs) from multiple cells.

The network consisted of a fixed number of observed neurons *N* while varying the total network size so that results could be compared across different levels of network observability. Network parameters were chosen to mimic ratios in mammalian cortex with neurons being 80% excitatory and 20% inhibitory^42^. (Supplemental Materials, S1: Neuron Model). These neurons were synaptically connected with uniform random probability. Neurons were injected with zero mean gaussian random current at each 0.5 millisecond timestep with variance tuned to elicit a 0.2 Hz average spontaneous firing rate per neuron across the network. These parameters were chosen based on reported spontaneous activity in Layer 2/3 vS1 barrel cortex and activity in layer 2/3 V1 in awake mice^26,37,43,44^. A subset of the network was randomly selected for observation to mimic experimental scenarios where only a subset of a large population is observable. A *T* x *N* binary random independent identically distributed stimulation matrix, M, was generated. Every 50 milliseconds a stimulation trial was carried out. This consisted of selecting a row without replacement from M and injecting current in neurons with nonzero indices in the row sufficient to reach firing thresholds. Voltage responses from each observable neuron in the network on the following time step were recorded for each trial. The full process of conducting a simulation run is summarized in the flow chart Figure 1B.

On top of this base model, we introduced other variables in the model that mimic realistic biological and technological factors. Specifically, synaptic failure was modeled as a binary random variable which sets the amount of current transferred from a pre-synaptic to post-synaptic neuron to zero. Off-target stimulation was modeled as instantaneous current delivered to a random single neuron that is not in the stimulation set in each trial with binary random probability of occurrence. Latency was modeled as a time delay between pre-synaptic action potential firing and evoked post-synaptic current (PSC). Specifically, for each connection in the network, latency takes on a value given by a normally distributed random variable rounded to the nearest time step. In models with random latency, voltage responses were measured in a three standard deviation interval following stimulation. Voltage decay for each cell was estimated by taking the mean of the v(t+1)/v(t) ratios over all post stimulation intervals. Responses for each time step over these intervals were normalized by subtracting the estimated decay then summed to obtain the total response for the respective trial.

Local cortical circuits may diverge from uniform sparsity and contain dense clusters of connections^45^ which could degrade mapping performance. The effect of clustered connectivity was explored by implementing a Wattz-Strogatz small world structure^46^. The connectivity matrix was constructed by first forming a ring lattice with *n* nodes and *k* edges depending on sparsity level. A third parameter, beta, controlled the probability of edges’ random reassignment. Beta spanned the range 0-1, with 0 generating a ring lattice of clustered local connectivity and 1 generating a random graph. After the digraph was generated, neurons were assigned to nodes while weights respective of cell types were assigned to edges.

### Decoding

The limits of compressive sensing were evaluated by varying the model parameters and quantifying map reconstruction using a confusion matrix. Where possible, parameters spanned the range from successful reconstruction to where no reconstruction was possible in fewer trials than the number of neurons. Values for network parameters are based on data from published experiments (Table 1). Each parameter studied spanned its range while other parameters were held at a base value.

**Table 1.** Parameter ranges tested. Each parameter spanned the indicated range while other parameters were held at the value in parenthesis.

Experiment parameters (base value)
|  |  |
| --- | --- |
| **Total Neuron in Network** | 200 – 4000 (1000) |
| **Observed Neurons** | 200 – 1000 (200) |
| **Sparsity** | 0.02 – 0.20 (0.1) |
| **Fraction of neurons stimulated per trial** | 0.1 – 0.5 (0.1) |
| **Off target stimulation probability** | 0.0 – 0.5 (0.0) |
| **Synaptic failure probability** | 0.0 – 0.5 (0.0) |
| **Latency mean (ms)** | 0.3 – 4.8 (1.0) |
| **Latency standard deviation** | 0.15 – 2.4 (0.0) |
| **Small world beta** | 0.0 – 1.0 (1.0) |
| **$\lambda$, sparsity-error trade-off** | (0.25) |

The basis pursuit solution (1) was solved for each measured neuron in the network using the CVX modeling system for convex optimization in MATLAB. A connection was declared to exist if a decoded weight exceeded a threshold relative to the strongest connection observed. An unconstrained version of equation (1) proved easier to solve in practice (Appendix 1.2.1):

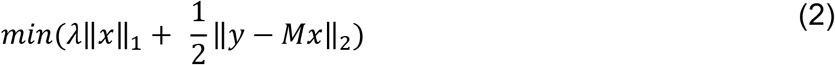

Here *λ* acts as a trade-off parameter between the sparsity of the solution and fit of the linear reconstruction. Entries in x greater than 1% of the largest weight were declared as true connections.

The objective function was further modified to include knowledge of cell type, which is typically known in cell-type specific optogenetic experiments. This was implemented to mitigate the limitation that not all the reconstructed post-synaptic weights were of the same sign. Neurons were classified as excitatory or inhibitory from the initial reconstruction results then solving the optimization with an additional set of constraints of the form:

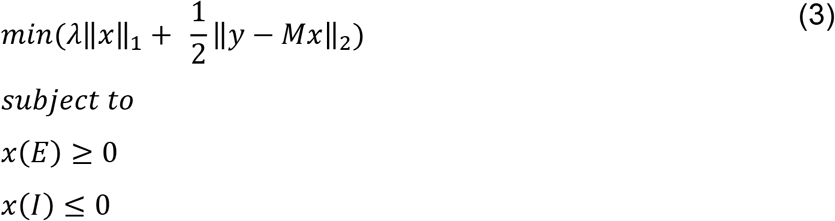

Where E and I are the indices of the excitatory and inhibitory neurons respectively.

## Results

The quality of the reconstruction was evaluated by comparing the estimated connectivity versus the ground truth. Connections correctly determined to exist or not exist were denoted true positives (TP) and true negatives (TN), respectively. Error types were defined as follows: Type I error, or false positive (FP), occurred when a connection was declared to exist in the reconstruction but did not exist in the actual model; Type II error, or false negative (FN), occurred when a connection was declared not to exist in the reconstruction but did exist in the actual model. As true positives are of greater interest than true negatives for this problem, recall and precision were used as metrics for comparison

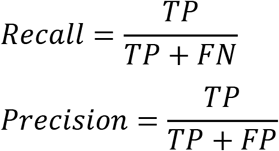

At 10% connection probability, we found that CS achieved a > 90% recall with only half the trials needed to achieve similar performance using single cell perturbation (Figure 2A). Similar results were recapitulated at each cell in the network, albeit with higher variance (Supplementary Materials, Figure S2). Higher recall was achieved in fewer trials in sparser networks except for the 2% connection probability case. Because the same λ was used for all cases, the sparsity was underestimated in this case resulting in excessive false positives. CS outperformed single cell stimulation until the probability of neuronal connectivity rose above 0.16 then performance fell off (Figure 2A). This drop in performance could be explained by the fact that compressive sensing reconstruction generally decreases with decreasing sparsity^47^. However, due to the partially observed, dynamical nature of the simulated neuronal network, changes in sparsity can contribute to errors beyond those suggested by CS alone. The amount of noise in the system is a function of the background firing, *e_n_sf__* (Figure 2D), which increases in more densely interconnected networks because the relative contribution of stimulated pre-synaptic currents becomes small compared to the total network activity in a given trial. Furthermore, as the network becomes more densely connected, more pre-synaptic inputs are being integrated at their respective post-synaptic cells. This creates larger fluctuations in currents which perturb the membrane voltage significantly thereby increasing *e_n_ν__* via the nonlinearities in the Izhikevich model^41^ (Supplementary Materials, Figure S1).

**Figure 2.**
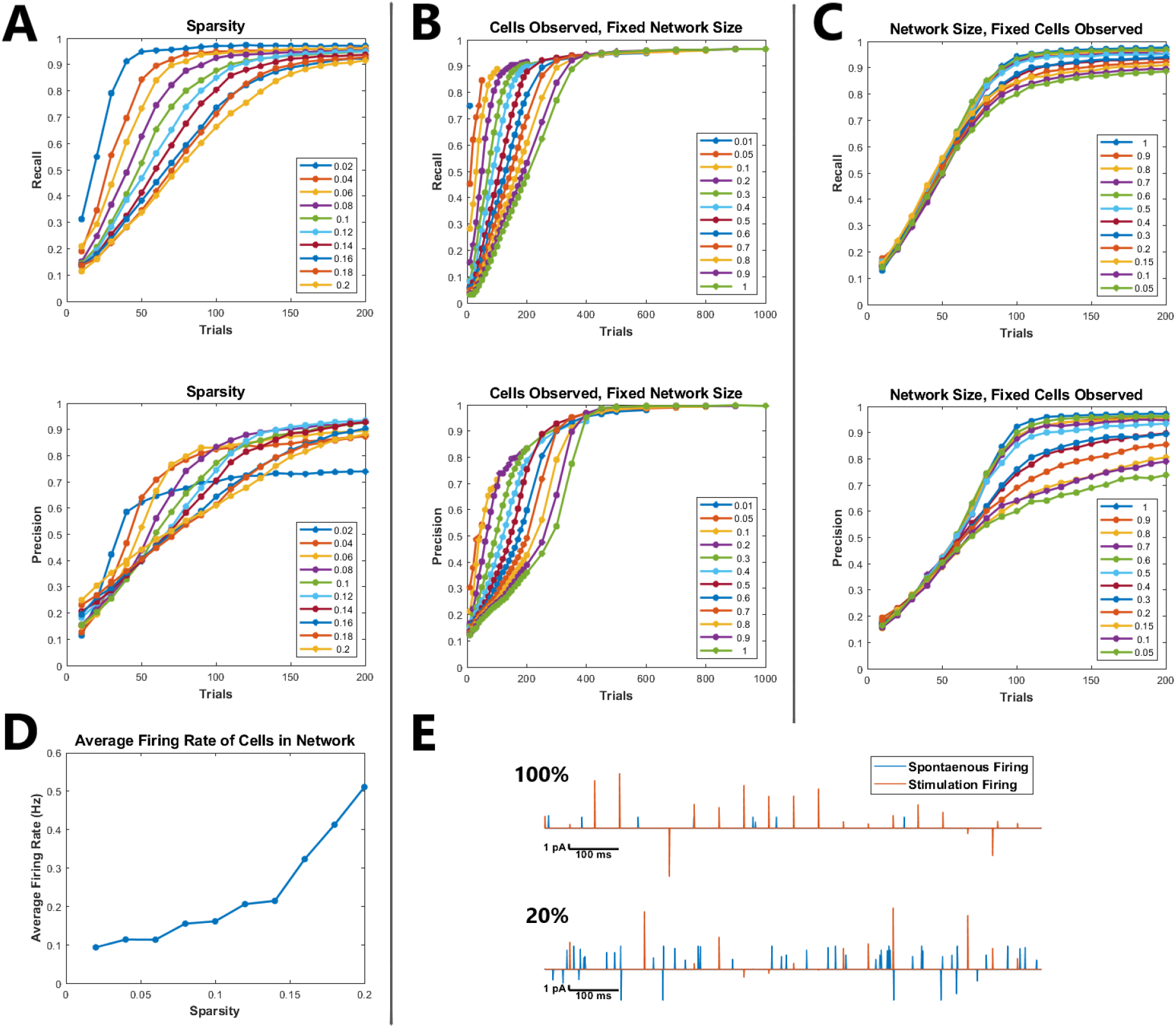
Performance as a function of network size, sparsity and observability. A. Precision and Recall performance as a function of sparsity. Each curve corresponds to a network with a specific probability of connection. B. Precision and Recall performance as a function of the observed population size. The size of the network size was held fixed at 1000 cells while the observed ensemble size was varied from a fraction of 0.01 (10 cells) to 1.0 (1000 cells). T = N trials were performed for each trace. C. Precision and Recall performance as a function of the overall network size. The size of the observed population was held fixed at 200 cells while the total network size was varied from 200 to 4000 cells. D. Average firing rates for cells in the network. Increasing connection probability increases background activity E. Examples of post-synaptic current events caused by stimulation evoked firing and spontaneous firing for a fully observed and partially (20%) observed network over a one second time interval.

For a fixed network size, we asked what effect the number of observable neurons have on reconstruction performance. Despite the decoding problem growing in complexity as more cells are added, we found that the recall and precision converge to the same points (Figure 2B). We then varied the network size as we held the number of observed neurons fixed. We found that performance dropped as the percentage of neurons observed decreased (Figure 2C). As the percentage of neurons observed decreased, the system became partially observed and spontaneous firing *e_n_sf__* began contributing significantly to the measured response (Figure 2E). This suggests that compressive sensing may be particularly well suited for models in which observing the entire network is possible such as C. Elegans^48^ or Zebrafish^49^.

We then asked whether varying the fraction of neurons simultaneously stimulated as part of a pattern affected the performance. We found that increasing this number beyond 20% of the observed ensemble in a given trial did not lead to substantial gain in performance (Figure 3A Top). This is in contrast to coherence metrics reported in the compressive sensing literature which suggest performance should increase with increasing nonzero entries in the stimulation matrix^50^. This deviation could be explained by the dependence between the size of the measured and stimulated sets of neurons. Whenever we stimulate a neuron in a trial, we cannot reliably measure the subthreshold response of that neuron as its depolarization causes a suprathreshold change in membrane conductance. As such, any postsynaptic current it might have received from other stimulated neurons in the same trial would be masked by the supra threshold response from that neuron leading to increases in *e_n_ν__* and consequently performance decline.

**Figure 3.**
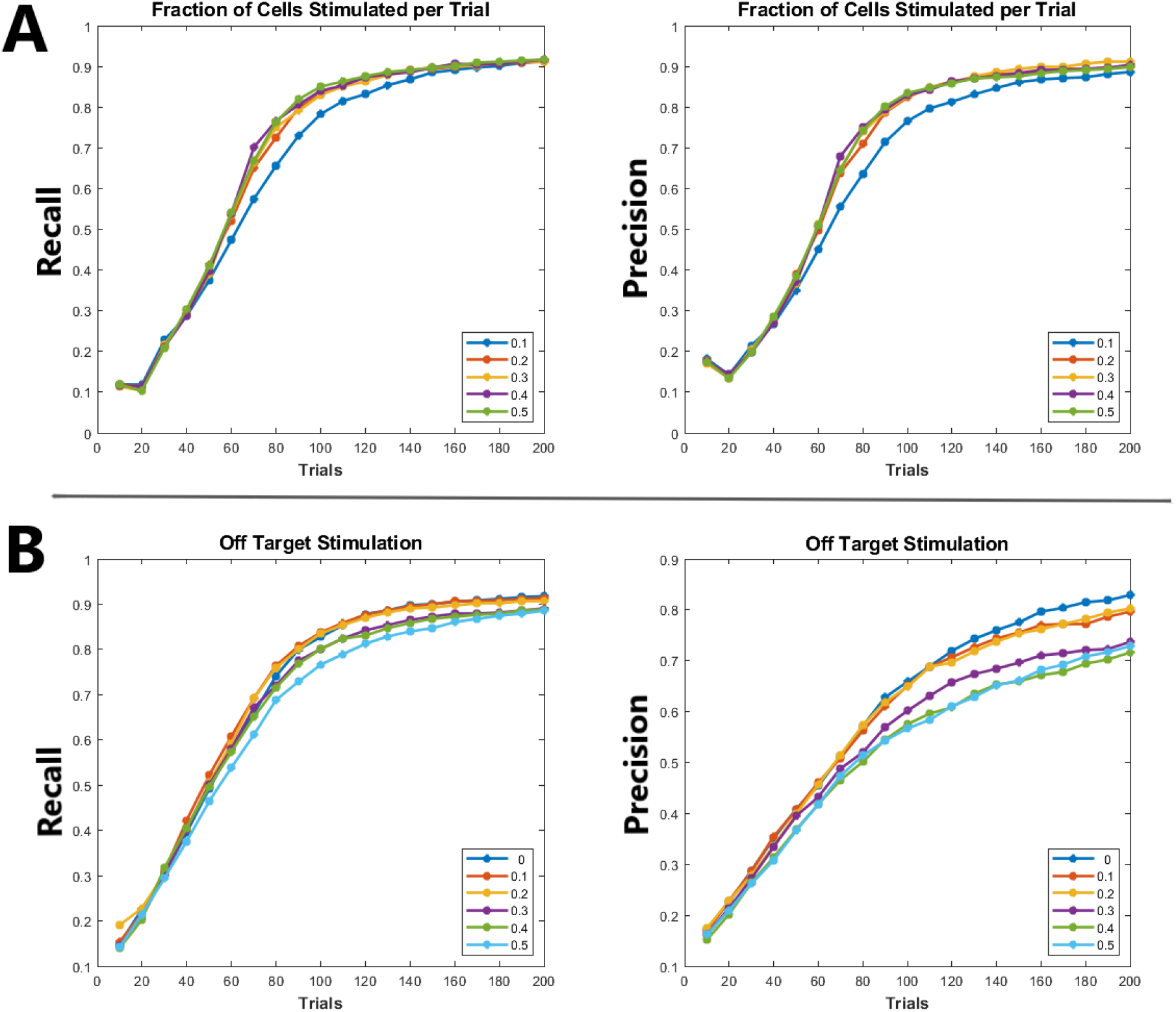
Performance under varying number of stimulated cells per pattern and off target stimulation. A. Precision and recall as a function of neurons stimulated per pattern. Each trace corresponds to the percentage of cells stimulated per trial in the observed population. B. Precision and recall as a function of off-target stimulation. Each trace corresponds to the probability that an additional neuron per trial was stimulated but not included in the reconstruction.

We then asked whether Off target stimulation had any effect on performance. We found the effect to be marginal (Figure 3B), likely due to the small size and random selection of the off-target neurons relative to the total being stimulated in a pattern. In practice, adequate characterization of laser illumination (e.g. when using computer generated holography combined with temporal focusing^51^) could limit this effect with knowledge of soma positions in 3D tissue volumes. Furthermore, simultaneous imaging through GECIs and GEVIs^16^ could ameliorate this issue by indicating which neurons fired concurrently with the directly targeted ones. The measurement matrix could then be adjusted post-hoc, placing ‘1’s in trials where cells fired and ‘0’s where they did not, to facilitate more accurate reconstruction.

We then asked whether synaptic failure had any effect on performance. This also mimics a Multi-Input Single-Output (MISO) system scenario in which only opsins are expressed in a given population while a single postsynaptic cell membrane response is measured at any given time (e.g. using whole-cell patch). In this case, variables such as opsin kinetics, expression level and laser power could lower the probability of spiking of presynaptic cells (despite the presence of highly reliable synaptic connectivity to the measured cell). We found that recall probability fell dramatically as the probability of synaptic failure increased (Figure 4A). Recall rates of 80% could be achieved but at high failure rates of ~50% in reconstruction. This resulted from solving an overdetermined system after 1000 trials for the 200 observed neurons. While neurons with these rates of synaptic failure have been observed in-vitro^52^, the mean total network rate is on the order of 15% which suggests that this worst case simulation results may not be indicative of all *in vivo* conditions. If high rates of synaptic failure are encountered, an alternative strategy could be to use cell type information during stimulation and limit the analysis to specific cell types. In fact, we found that by limiting the observed population to only excitatory and constraining the reconstruction to specific cell types resulted in a substantial increase in performance (Figure 4B).

**Figure 4.**
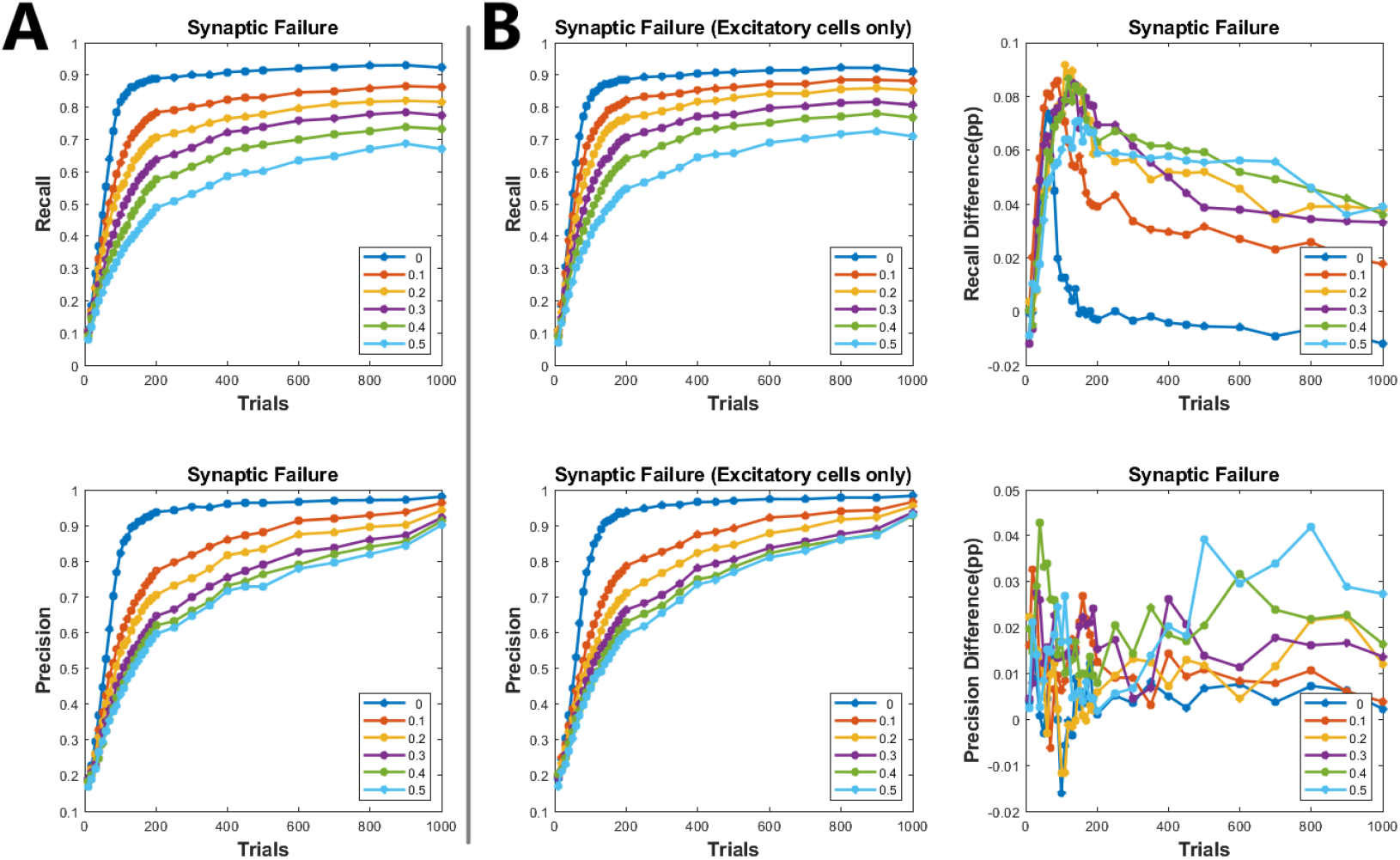
Performance for unreliable synaptic transmission, which also mimics failure to fire a presynaptic neuron, for example, by insufficient laser power/slow opsin kinetics or reduced cell-specific opsin expression. A. Precision and recall as a function of synaptic failure. Each trace corresponds to the probability that an action potential in a pre-synaptic cell did not propagate to the post-synaptic site. B. Left: Precision and recall as in A but for an observed network of only excitatory cells. Right: Difference between the two reconstructions in A and B. Percentage change in recall and precision in the observed excitatory network.

We then asked whether performance was affected by post-synaptic response timing, which could be a function of cell refractoriness, opsin kinetics and overall brain state. We found that recall probability remained robust as the mean and standard deviation of response latency increased (Figure 5). However, precision declined for mean latencies above 2.4ms and standard deviations above 1.2ms. As the mean and standard deviation of latency increase, the interval over which responses must be summed increased. This spreading of the response over a longer interval not only decreases the amplitude of responses relative to the background noise but introduces the possibility of firing from higher order projections outside of the observed set (polysynaptic effects). However, other studies suggest this type of noise can be tolerated as *in vitro* data for rat pyramidal visual cortex cells had latencies well within the range used here for reconstruction^53^.

**Figure 5.**
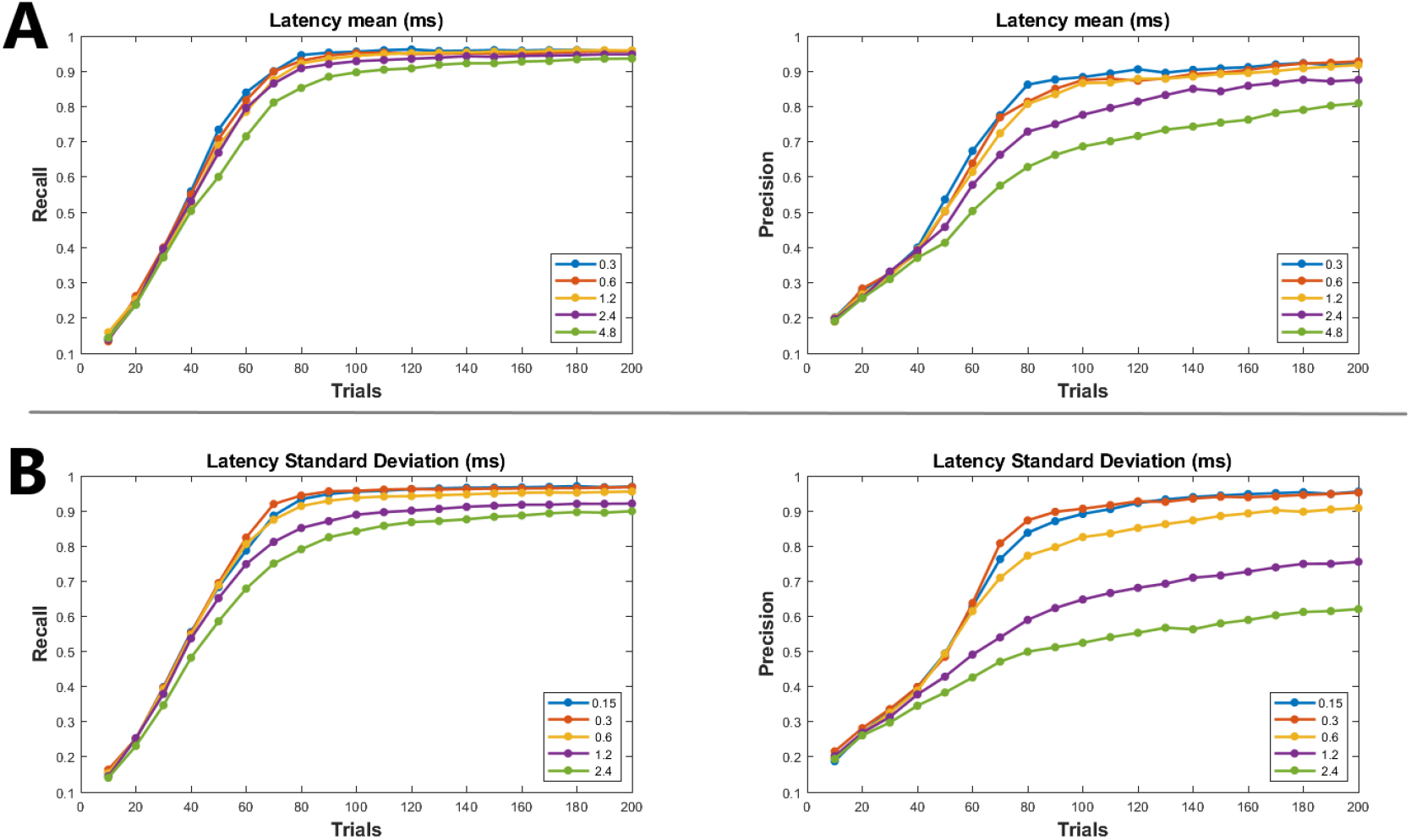
Performance against variable latency. The propagation delay for each connection was drawn from a normally distributed random variable with fixed mean and variance. Mean and standard deviation were independently varied while the other variable was held constant. A. Precision and recall for a fixed 0.6ms standard deviation with each trace corresponding to a different mean. B. Precision and recall for a fixed 1.2ms mean with each trace corresponding to a different standard deviation.

Single cell simulations (Supplemental Figure: S2) also showed good recovery with parameters similar to this data. We last asked whether network topology had any effect on performance. We found that recall probability remained robust despite varying the small world clustering characteristics of the network (Figure 6). This may have been a result of the regularity of the degree distribution of the Wattz-Strogatz model where compact support^34^ can be easily satisfied. Models with high variances in degree distributions, such as scale-free networks^54^, which strain the sparsity requirement of compressive sensing may result in poorer performance.

**Figure 6.**
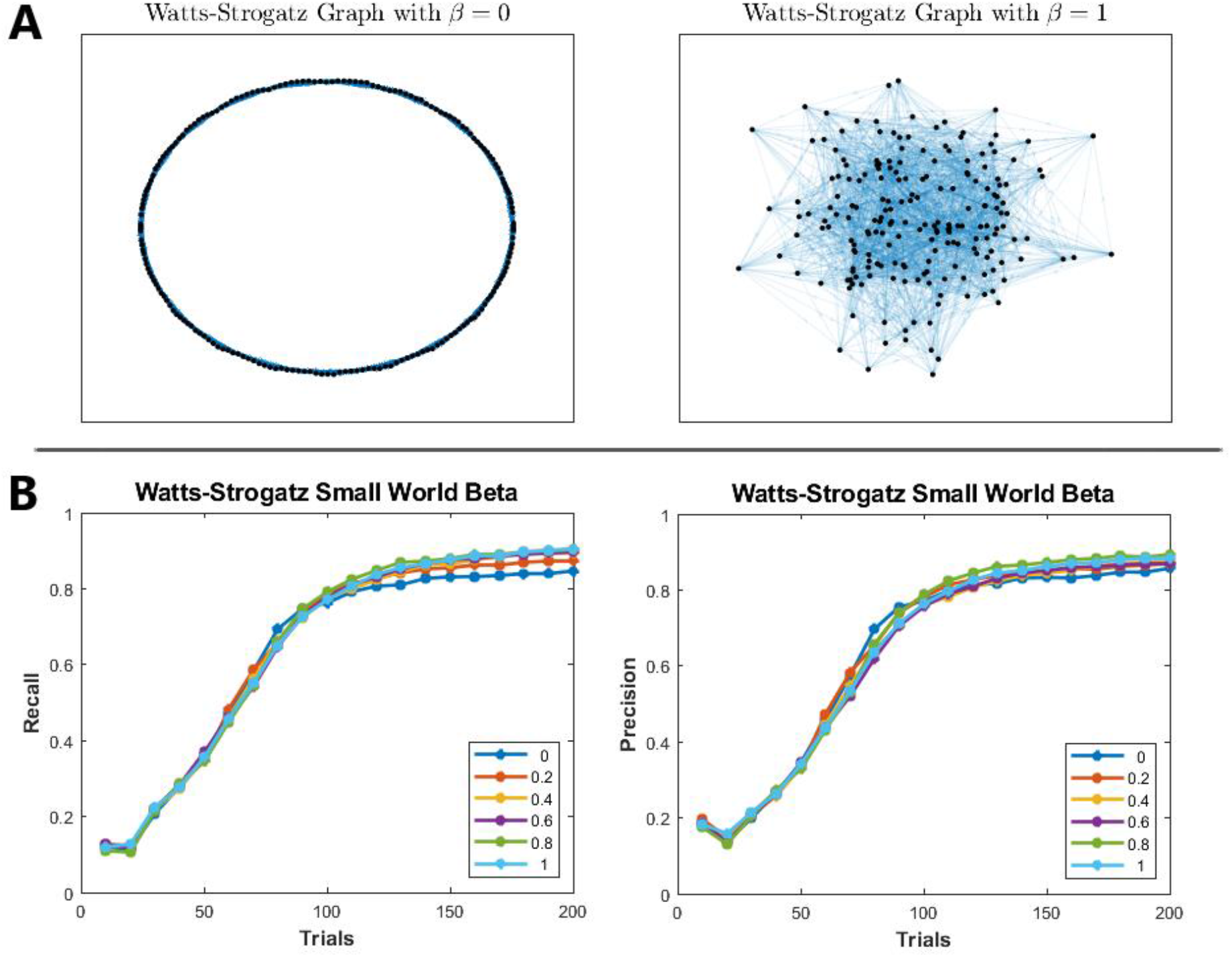
Performance for small-world network topologies. A. Illustrations of Wattz-Strogatz graphs for beta values of 0 and 1 where 0 represents a ring lattice structure with highly local connectivity (left) and 1 represents a randomly connected network (right). B. Recall and precision for small world beta. Each trace corresponds to networks with a different beta parameter.

## Discussion

In this work, we explored the use of parallel stimulation and compressive sensing for mapping synaptic connectivity between neurons with single cell resolution, cell type specificity and nonlinear dynamics. We constructed a large network model with parameters derived from biological experiments to quantify multiple aspects of the performance. At predetermined time points, subsets of neurons were stimulated and PSP responses immediately following the stimulation intervals were measured for each cell. These responses were subsequently decoded by solving a constrained linear programming problem that estimates each presynaptic neuron weight. For each set of parameters tested, recall and precision were computed for the estimated synaptic weights and compared to the ground truth weights. Results demonstrate that parallel stimulation and compressive sensing can be synergistically used to map connectivity over a wide range of biologically plausible parameters in far fewer trials than sequential stimulation methods. The wide range of parameters tested also provided further insight into the technique’s limitations.

Prior work has applied compressive sensing to the synaptic connectivity mapping problem, albeit in highly simplified simulation settings or from postmortem anatomical data ^36,55,56^. More recent work has applied this approach combined with parallel stimulation of visually identified cells with measured single cell PSCs *in vitro*^38^. Here, we pushed the limits of the simulation environment by incorporating much larger network models with recurrent connectivity and varying many critical parameters to closely resemble biological networks. We chose a membrane voltage-based model for ease of implementation and to mimic a voltage imaging experiment where PSPs from multiple cells are recorded simultaneously. PSCs from whole cell recordings can be used just as readily as PSPs, by substituting their peak amplitudes into y of equation (1), as both are measurements proportional to synaptic strength. Our results suggest that the idea of parallel stimulation and compressive sensing is a useful method for high resolution connectivity mapping under different scenarios of network sparsity, observability, reliability, and latency. We found that that network sparsity and synaptic reliability were primary determinants of the performance. Unreliable synapses greatly degraded performance, particularly when cell type specific information was not used. Imposing constraints on the accepted solutions considering individual cell types improved the performance. Furthermore, increasing the number of cells stimulated per trial improved reconstruction but plateaued when the average network firing rate reached a certain level that eventually masked the measured PSPs in the decoding step.

The limitations of the modeling framework in our study must be considered. First, while the partially observable network model recapitulated biological processes of presynaptic spiking probability/synaptic failure, random propagation latency and variable topology^57^, it was assumed that these parameters remained constant over the course of a given simulation run. In practice, optically evoked presynaptic spiking and synaptic reliability can be a function of many variables such as opsin expression level, time since last spike^53^ and up/down brain states^58^. Second, simulation parameters were varied independent of one another in a given simulation run. In practice, they might be interdependent, for example, in a synchronized cortical state^59^. Third, changes in membrane responses were governed by differential equations and constant noise processes in the model. Voltage indicator kinetics were not included in the model and it was assumed that the measured change in membrane voltage followed the model equations. This made the sum of the integrated PSPs, on average, to be directly proportional to peak voltage amplitude during the interval immediately following stimulation. In live experiments, however, a matched filtering approach might be required to weigh each time point in the post-synaptic events^60,61^. Fourth, off target stimulation might cause responses from proximal and apical dendrites from other cells which must be filtered out from the measurement^60^. These differences should be accounted for when applying the method *in vivo*.

Compressive sensing assumes that the measurements taken to perform reconstruction are linear superpositions of components within their respective measurement and therefore the largest performance decrease occurred when these assumptions were violated. Care must be taken when estimating the sparsity of the network as an untuned λ, the parameter which controls the tradeoff between the number of components in the solution versus the error, can lead to false positives if underestimated and false negatives if overestimated. In practice, underestimation may be preferred as false positives are much easier to check for with single cell stimulation than false negatives. Checking for false positives would require trials on the order of the number of connections which is much smaller than the number of total possible connections. Reconstructions that did not constrain neurons to either be excitatory or inhibitory suffered slightly in performance. Recall and precision fell precipitously as probability of presynaptic spiking/synaptic failure increased. While this was somewhat ameliorated by limiting the observed network to only excitatory cells and constraining the reconstruction, the applicability of CS based reconstruction will be limited in areas know to have high levels of unreliable synaptic propagation such as hippocampus^62^. All things considered, our proposed method remains highly promising for a wide range of neural circuits that obey the bounds of high fidelity reconstruction^23,25,26,28,29,43^. Future work could focus on adapting it to work more accurately in the presence of unreliable synapses and efficient sampling strategies with varying levels of sparsity.

## Supplemental Materials

### S1: Single Neuron Model

The neuron model used was the Izhikevich model.^41^ It can be described as a biologically plausible model which is computationally simple but capable of producing rich firing patterns for a variety of cell types. The evolution of the system is governed by a system of two-dimensional ordinary differential equations:

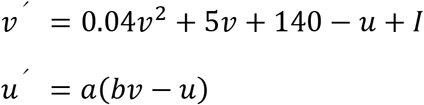

With the auxiliary after-spike resetting

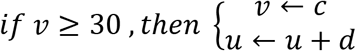

Where *u* and *ν* are dimensionless variables; *a, b, c* and *d* are dimensionless parameters; ‘ denotes the derivative with respect to time t. The variable *v* represents the membrane potential of the modeled neuron while *u* acts as a membrane recovery variable motivated by the biophysical process of potassium ion current activation and sodium ion current inactivation, giving negative feedback to *ν*. Following a spikes peak of 30mV, *u* and *ν* are reset. The variable *I* represents incoming synaptic or injected currents. It is calculated by summing the weights, *S_i,j_*, of the pre-synaptic neurons, *j*, to post-synaptic neurons, *i*, that fired in the previous timestep with a zero-mean, normally distributed random input with fixed cell-type specific variance P.

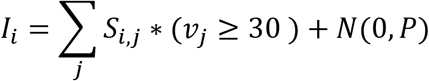

The parameter *a* describes the rate of recovery of *u* with smaller values resulting in slower recovery. The parameter *b* couples the membrane recovery *u* to the membrane potential *ν* with greater coupling leading to dynamics such as subthreshold oscillations and low-threshold spiking. The parameters *c* and *d* correspond to the reset values, following a spike, for the membrane potential *ν* and membrane recovery variable *u*, respectively. Different choices of parameters can lead to neuron behaviors which span what is seen in the brain. I chose to only include parameter choices which modeled behaviors observed in cortex (excluding thalamo-cortical neurons) as that would likely be the first in vivo area studied due to its proximity to the surface of the brain and the depth limit of multiphoton microscopy. Excitatory neurons span parameter ranges between RS (regular spiking), IB (intrinsically bursting) and CH (chattering) neurons. Inhibitory neurons span the parameter ranges between FS (fast spiking) and LTS (low-threshold spiking) neurons.

Relating these components in terms of CS measurements equation (1), the stimulated firing corresponds to *Mx* while the spontaneous firing and membrane dynamics form e. Let the superscript *c*, typically the set complement, denote neurons in the network model but not in the observed set, and define *M_K_* as a T x K matrix that takes value ‘1’ if a neuron in the respective column fired in the timestep prior to a measured response. We can then write the formula for y in terms of the model equations evaluated at the trial times as:

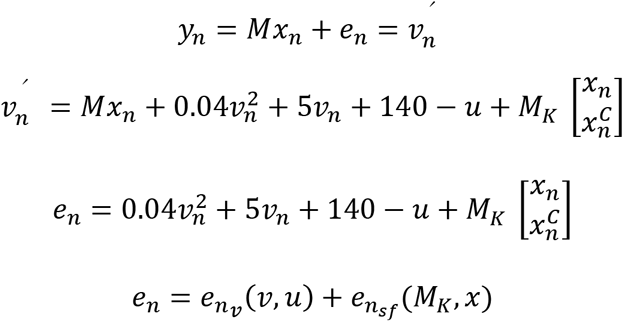

This divides the deviation from a linear summation into two terms: *e_n_ν__* which is dependent on local membrane states and dynamics, and *e_n_sf__* which is dependent on the spontaneous firing of neurons projecting onto the measured neuron.

**Figure S1.**
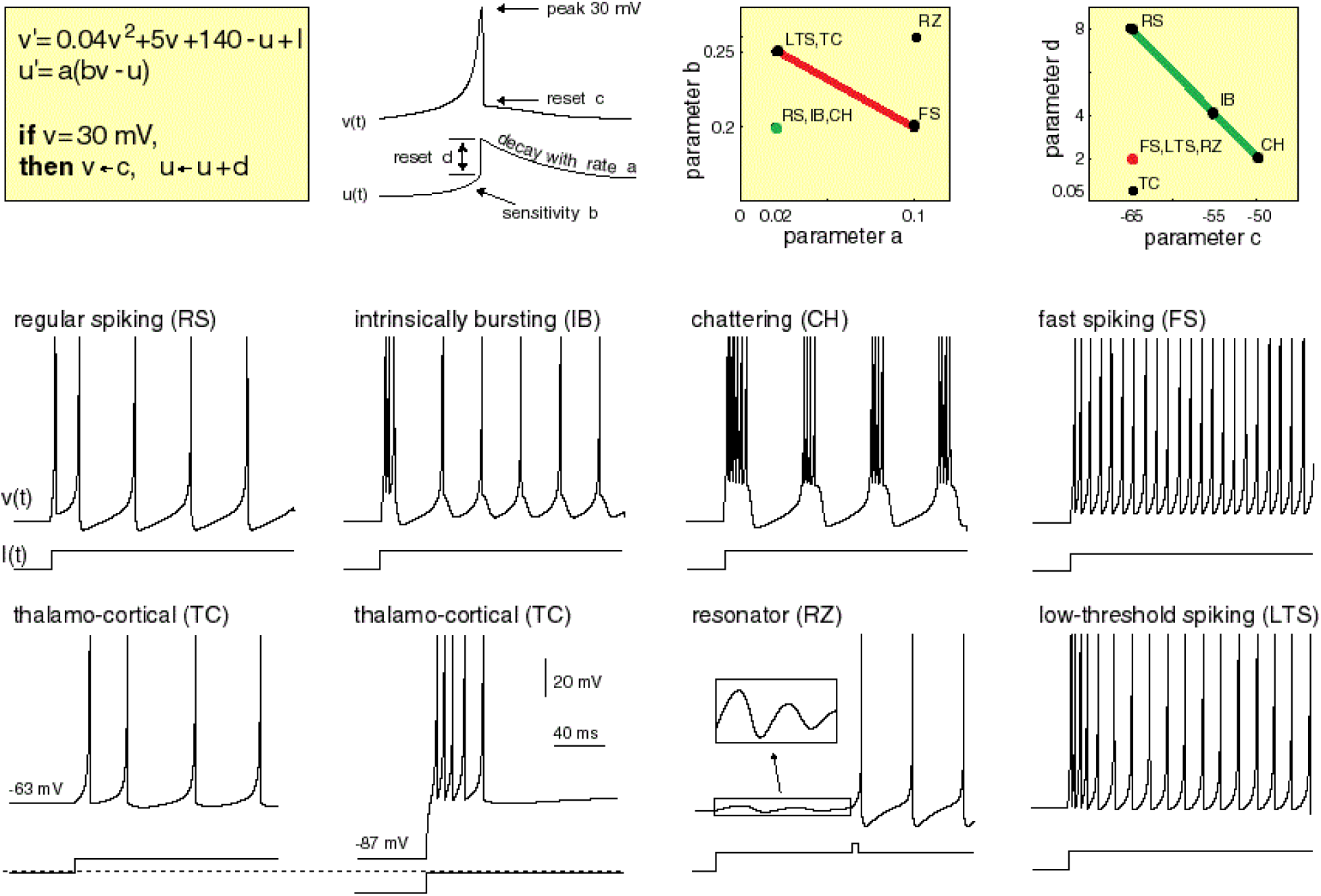
Izhikevich model neurons. Top insets correspond to model dynamics and parameter relationships. Bottom insets correspond to neuron behaviors for different parameter sets. Firing propagates through the model via the ‘I’ term where ‘I’ reflects the sum of the weighted connectivity of presynaptic neurons that fired in the previous timestep. Regular spiking neurons respond with a short inter-spike period when first presented with a stimulus then steadily increase in period for the length of the stimulus in a phenomenon known as spike frequency adaptation. Intrinsically bursting neurons fire a stereotypical burst of spikes when first presented with a stimulus followed by repetitive single spikes. Chattering neurons fire stereotypical bursts of closely interspaced spikes. Fast spiking neurons can fire high-frequency periodic trains of action potentials without any spike frequency adaptation. Low-threshold spiking neurons act like fast spiking neurons but have lower firing thresholds and display spike frequency adaptation. Parameter ranged used for excitatory neurons in this proposal are highlighted in green while ranges used for inhibitory neurons are highlighted in red. This modified figure was reproduced with permissions from the author.

### S2: Performance under randomized trial selection

In order to assess the effect of trial order on reconstruction performance, the experiment was repeated using randomly sub-sampled trials. The network was initialized with base values for all parameters (1000 Total neurons, 200 observed neurons, 0.1 sparsity, 10% observed neurons stimulated per trial, λ=0.25, propagation latency of 1ms). For a 10 neuron subset of the observed network, 100 simulations were performed for each level of subsampling. In each simulation, T trials were randomly selected, without replacement, from N total trials to be included in the reconstruction. The effect of trial order was measured using the mean, 10^th^ percentile and 90^th^ percentile of recall and precision at each level of subsampling (Figure S2). For every neuron, the reconstruction converged as more trials were included.

**Figure S2.**
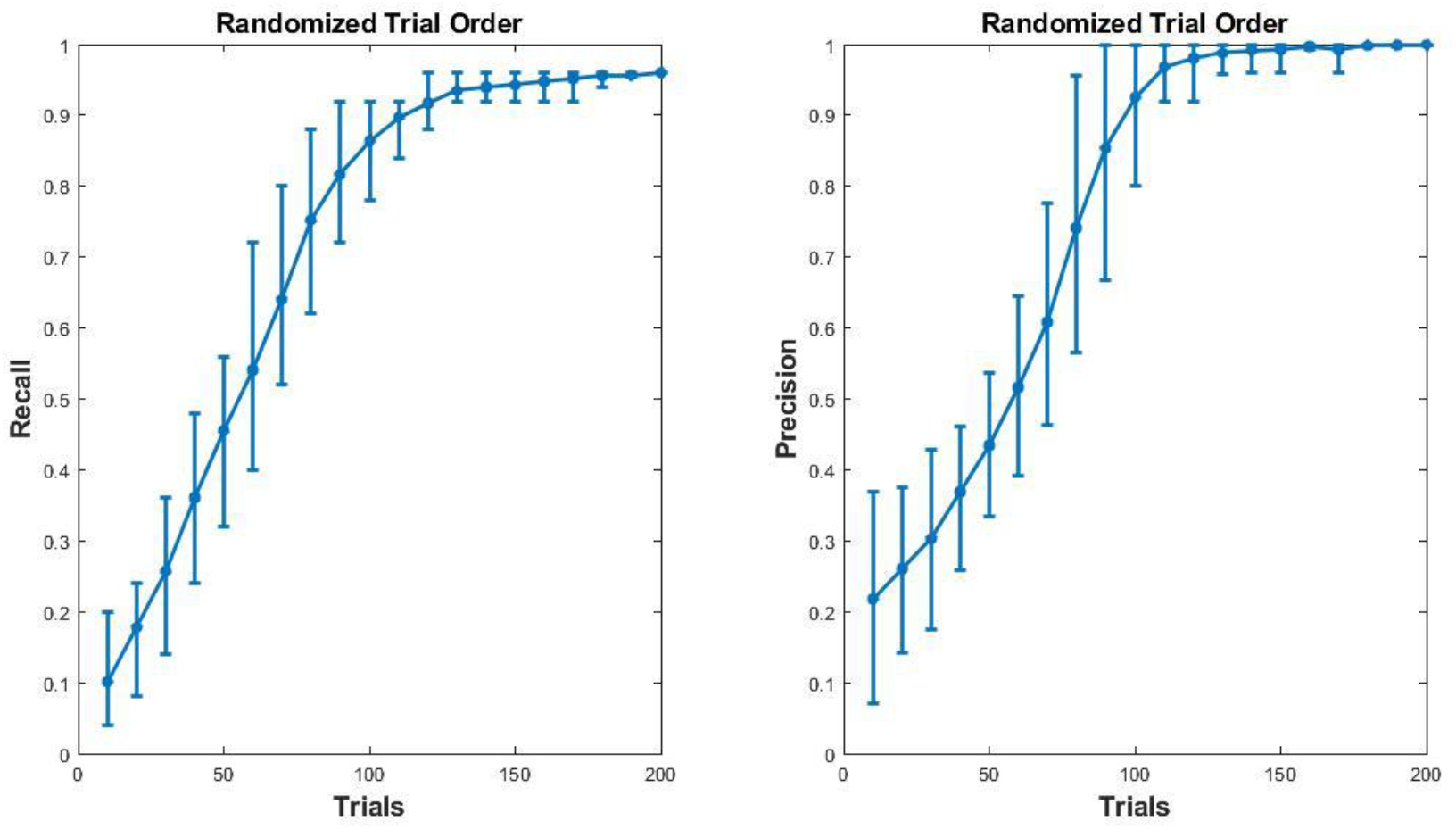
Single cell performance with random trial selection. Recall and precision for 100 randomly sub-sampled reconstructions for neuron 9. Traces correspond to mean performance while error bars span from the 90^th^ to 10^th^ percentile.

